# Clonal evolution and genome stability in a 2,500-year-old fungal individual

**DOI:** 10.1101/377234

**Authors:** James B. Anderson, Johann N. Bruhn, Dahlia Kasimer, Hao Wang, Nicolas Rodrigue, Myron L. Smith

## Abstract

In the late 1980s, a genetic individual of the fungus *Armillaria gallica* that extended over at least 37 hectares of forest floor and encompassed hundreds of tree root systems was discovered on the Upper Peninsula of Michigan. Based on observed growth rates, the individual was estimated to be at least 1500 years old with a mass of more than 10^5^ kg. Nearly three decades on, we returned to the site of individual for new sampling. We report here that the same genetic individual of *A. gallica* is still alive on its original site, but we estimated that it is older and larger than originally estimated, at least 2,500 years and 4 × 10^5^ kg, respectively. We also show that mutation has occurred within the somatic cells of the individual, reflecting its historical pattern of growth from a single point. The overall rate of mutation, however, was extremely low. The large individual of *A. gallica* has been remarkably resistant to genomic change as it has persisted in place.

## 1. Introduction

In the late 1980s, Smith, Bruhn, and Anderson [1] discovered a genetic individual (designated C1) of the fungus *Armillaria gallica* that extended over at least 37 hectares of forest floor and encompassed hundreds of tree root systems in Michigan’s Upper Penninsula. Based on observed growth rates, C1 was estimated at the time to be at least 1500 years old with a mass of more than 10^5^ kg. The conclusion was that C1 was among the largest and oldest organisms on earth, a remarkable claim given that Armillaria is essentially a microorganism existing largely as microscopic hyphae embedded in their substrates. Nearly three decades on, we returned to the site of C1 with the tools of whole-genome sequencing. Here, we show that C1 is still alive on its original site, but it is older and larger than originally estimated, at least 2,500 years and 4 × 10^5^ kg, respectively. We also show that mutation has occurred within the somatic cells of C1, reflecting its historical pattern of growth from a single point. The overall rate of mutation, however, was extremely low. On the spectrum of mutability in somatic cells, Armillaria occupies the extreme of stability, opposite the extreme of instability as typified by cancer [2].

Since the discovery of the C1 [1, 3] individual, much has been learned about genomes, gene content, and gene expression in Armillaria. A general picture of how individuals of this fungus become established and persist in woodlands has emerged. Because of its broad host range, capacity for enzyme secretion, and prolific production of rhizomorphs, its unique organs of local dispersal, Armillaria appears to be adapted for persistent growth in local habitats [4]. Since its discovery, it has become clear that C1 is not unique in its size and age. Any temporally continuous forest could support large, old Armillaria individuals. Indeed, at least two other individuals of a sibling Armillaria species (*A. solidipes*) have been reported to occupy larger areas than C1 [5–7].

Armillaria lives both as a saprophyte on dead wood and as a necrotrophic parasite, killing host plant tissues in advance of its growing mycelia by secreting hydrolytic enzymes and possibly virulence factors [8]. The range of potential host plants is broad, encompassing both angiosperm and conifer trees [9]. The large genome size of Armillaria relative to its sister taxa is due to increased gene content, particularly in plant cell wall degrading enzymes, rather than to the proliferation of transposable elements [8]. Individuals of Armillaria are established when mating occurs between haploid spores or their germlings [10, 11]. After mating, a diploid mycelium is established rather than the dikaryotic mycelium typical of other basidiomycetes. From mycelium, rhizomorphs develop and push through the soil and decaying woody substrates in much the same way as a plant root. Rhizomorphs function as foraging organs in locating new sources of food in dead wood or weakened trees. They can also persist in stasis in soil until new substrates become available. Gene expression in rhizomorph development shares much the same signature of multicellularity found in fruit body development [8]. Rhizomorphs also share with vegetative mycelium a pattern of gene expression supporting enzyme production [8]. Against this background, the focus of this study was on the mutability of Armillaria - detecting genomic change in the spatial record left from the proliferation of somatic cells in space from a single zygote ancestor.

## 2. Methods summary

### (a) Sample

In 2015-2017, we revisited the location of C1 and made 245 collections linked to GPS coordinates (Table S1). Collections were mostly pure cultures from rhizomorphs, but in the fall of 2015 and of 2016, samples also included fruit-body tissues which were used directly in DNA analysis. Subsequent methods generally followed Anderson and Catona [12].

### (b) Culturing and DNA extraction

Rhizomorphs were cut into 2 cm segments and placed in 2.5% hypochlorite bleach for ten minutes to surface disinfect. The rhizomorphs were then trimmed to less than 1 cm and placed on 2% malt-extract agar medium. Liquid cultures were in 2% malt extract without agar. Mycelium was harvested, flash frozen in liquid N2 and then lyophilized. DNA was extracted with a CTAB-low salt CTAB precipitation method as described earlier [13].

### (c) Somatic compatibility testing

To determine whether or not a collection belonged to C1, we tested for somatic compatibility (Table S1). Somatic compatibility distinguishes mycelia of the same individual which merge seamlessly in culture from mycelia of different individuals which react with a zone of cell death and pigmentation [14]. We noted one large grouping of 110 isolates that were later confirmed to represent C1 and a smaller grouping of eight isolates that matched the C2 described earlier [1, 3]. C1 and C2 are still centred on the same respective localities as reported in 1992 [8].

### (d) Initial genotyping

Polymorphic molecular markers were also used to test whether a new collection represented the C1 identity or not (Table S1). For example, in one segment of DNA, homozygosity for absence of a MboI site has a frequency of 0.64 in the general population [15]. In addition, the 3’ end of the 25S rRNA gene is heterozygous in C1 for a length polymorphism, a genotype that has a frequency of 0.21 in the population. The frequency of the combined genotype of the C1 individual over the two DNA regions is 0.13. The combination of the two DNA regions and somatic compatibility drives the probability of a spurious match with a non-C1 individual much lower.

### (e) Illumina sequencing

HiSeq Illumina sequencing was by paired-end with 155 bp reads at the Centre for Applied Genomics at the Hospital for Sick Children, Toronto. The Illumina sequences for 15 collections of C1 are deposited as accession PRJNA393342 in the SRA at NCBI.

### (f) Bioinformatics

The pipeline mapped the raw fasta files onto the reference genome, produced the pileup files from the resulting.bam files, and then filtered the pileup files to discover the variation among the Illumina sequenced strains. The pipeline is described in detail in Bioinformatics S1.

## 3. Results and Discussion

The C1 individual was first identified in the late 1980s by making spatially mapped collections of the fungus on the Michigan site and then genotyping them over multiple loci[1, 3]. The C1 designation used here corresponds to Clone 1 in the original publication [1] and to “The Humungous Fungus” as named by the news media. All collections of C1 had the same multilocus genotype and shared an identical mitochondrial type. Other nearby individuals had different multilocus genotypes and mitochondrial types [3, 15]. Another individual of lesser spatial extent, designated C2 here, was described in addition to C1.

In 2015-2017, we revisited the site of C1 and made 245 collections linked to GPS coordinates (Table S1). Figure 1 shows distribution of isolates representing C1, C2, and all other genotypes (which were excluded here from further analysis). Fifteen of these collections of C1 were Illumina sequenced (Table S1) to approximately 100X average coverage (NCBI Accession: PRJNA393342). The sequence reads were initially aligned with a 98 kb mtDNA reference, which was derived from another individual of *A. gallica* from Ontario, Canada [12] (JGI *Armillaria gallica* 21-2 v1). C1, C2, and three additional individuals from the Michigan site each had a unique mtDNA genotype based on well-defined SNPs (Table S2). The recent Illumina sequenced collections of C1 have the same mtDNA genotype as a living strain of C1 that was collected in the late 1980s. Similarly, the recent collections of C2 are linked to a C2 strain from the late 1980s by having the identical mtDNA genotype (Table S2).

**Figure 1.**
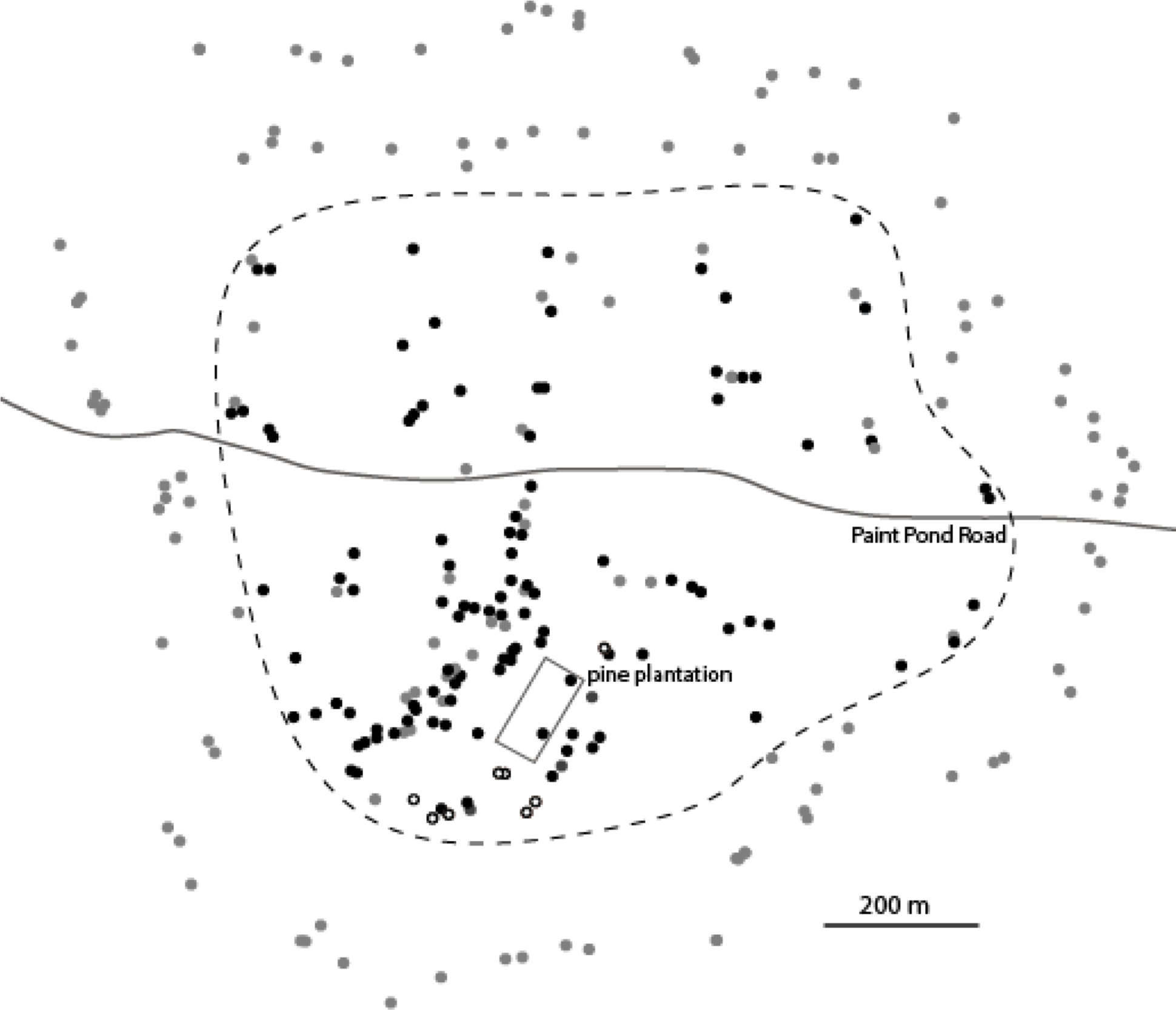
Map of all collections of Armillaria. Black dots, C1; open circles, C2; gray dots, all other individuals combined. The outline of the pine plantation and Paint Pond Road are included as alignment features. The dashed line encompasses collections of C1 (and includes some non C1 collections. The present sample, which was larger and more broadly distributed than the previous[1], was designed to find the approximate borders of C1. The present sample reveals that C1 is larger and older than originally reported[1]. Based on previous growth rate measurements and estimation of fungal biomass, the revised estimates for minimum age and mass are 2,500 years and 4 × 10^5^ kg, respectively.

To test for the signature of mutation, we searched for variation in the nuclear genome of the Illumina sequenced strains. After all filters were applied, 163 variants were found (Table S3). In this search, the key requirements for eliminating false calls were to require (a) that candidate sites had at most two alleles among the sequence reads, one reference and one alternate, (b) that one or more strains have 0% or 100% of the reads representing the alternate allele (a “purity” criterion), and (c) that in the heterozygous sites the allele frequencies hovered around 50% (30-70%).

Two general kinds of genetic changes were seen. First, for 151 cases, only single sites experienced the change and adjacent sites in the genome were not affected. We interpret these changes as point mutations resulting in a gain of heterozygosity. The heterozygous strains after mutation had a nearly equal mix of reference and alternate alleles among the Illumina reads, while the homozygous strains had purely one allele. Second, there were regions in strains that were homozygous at a number of adjacent sites that were otherwise heterozygous in other strains; we interpret these events as loss of heterozygosity (LOH) in which a string of adjacent SNP sites that were formerly heterozygous all become homozygous simultaneously, as can happen with mitotic gene conversion. That only six LOH tracts were observed (Table S3), and that the preponderance of the original heterozygosity in the individual has been maintained, is remarkable given the potential for mitotic gene conversion and crossing over, which would lead to homozygosity over substantial portions of chromosome arms.

The predominant pattern of both point mutation and LOH was that genetic changes were observed as singletons, present in no more than one Illumina sequenced strain (130 of 163 sites, Table S3). As expected for recent mutations mostly unaffected by selection [16], the majority (73%) of point mutations were due to C to T transitions (Table S3).

On average, each strain had about 10 singleton changes that were not shared among the other strains. A minority of changes (33 sites), however, were shared among two or more of the strains. These shared changes were particularly informative because they reflect the historical growth pattern of the individual, while the singletons reflect a history of units of the individual persisting in place over time, presumably after the period of general expansion had occurred. This interpretation arises from the fact that individuals of Armillaria cannot exist in stasis over the long term. Existing food sources become exhausted and new sources of food become available, meaning that growth of the fungus is necessary even to remain in the same place.

To examine the spatial relationships of the mutations, we constructed a phylogeny of 14 Illumina sequenced isolates of C1) using the variants in Table S3 as characters and maximum parsimony as the optimality criterion (Fig. 2). This approach was feasible because a branching filamentous fungus, whether growing in a petri dish or on the forest floor, represents a microcosm for the bifurcating tree of life. The phylogeny shows a high degree of internal consistency and the nesting of the clades in space can be interpreted as reflecting past growth and colonization patterns. Three of the 33 sites, however, showed homoplasy in the phylogeny; at these sites, changes occurred two or three times in different branches of the tree. These examples are unlikely due to independent mutation. With a low mutation rate and approximately 100 mb in the genome, the candidate sites for mutation are essentially infinite. The alternative explanation is that the parallel changes could represent single somatic recombination events during the early expansion phase after the individual was established, with the recombinants then becoming spatially separated within the individual. The potential of somatic recombination in fungi has ample precedent [17, 18].

**Figure 2.**
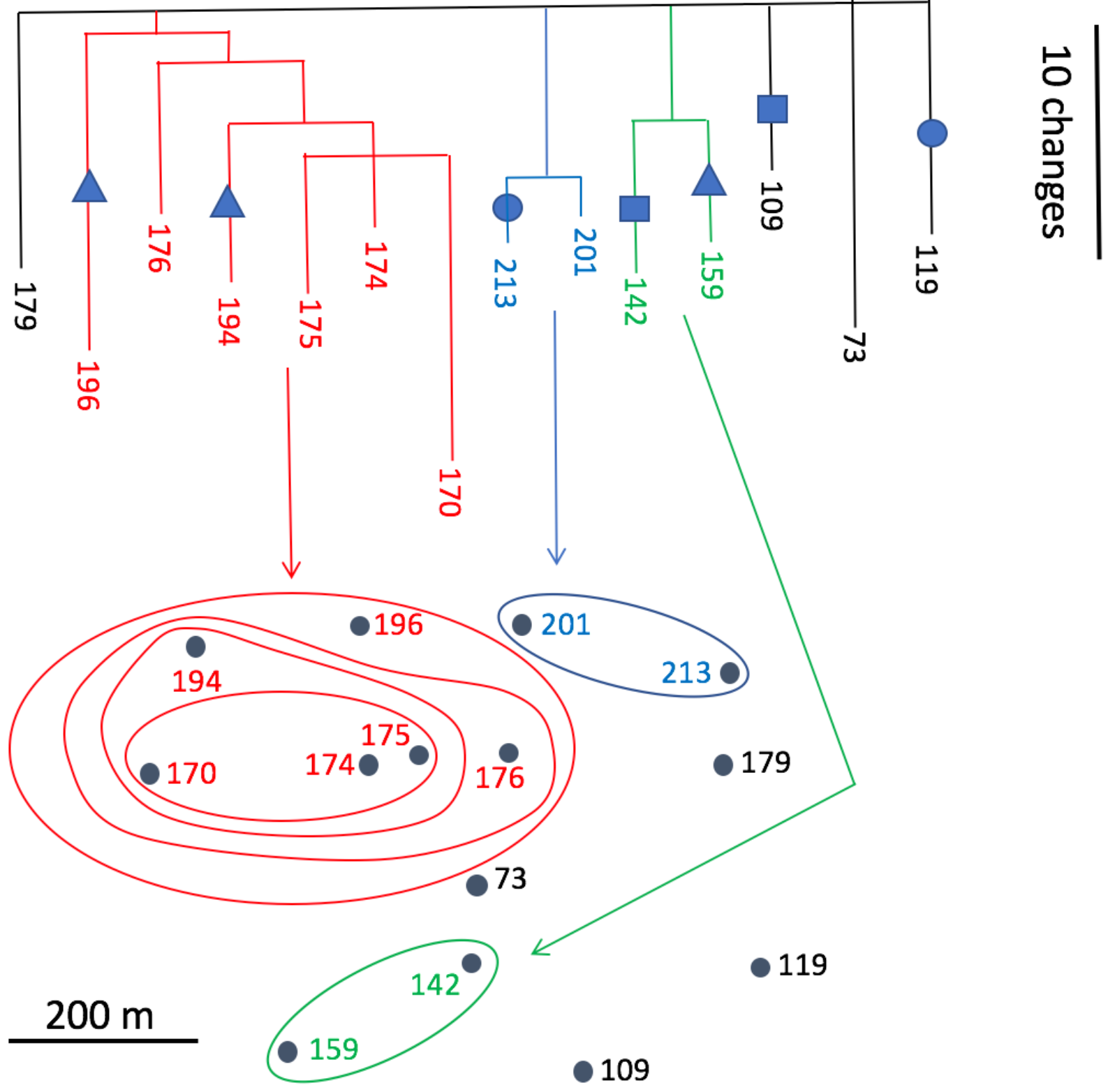
Phylogeny of Illumina sequenced strains of *A. gallica* tied to spatial origin. The variants for this geophylogeny are listed in Table S3. Symbols represent sites at which changes occur in different parts of the tree due either to recurrent mutation (unlikely) or recombination. Circles, contig 12, position 174630; squares, contig 61, position 336513, triangles, contig 33, position 701686 (Table S3).

Next, for an independent test of the phylogenetic pattern among Illumina sequenced strains, we tested nine sites in Table S3 among all isolates of C1. Once again, the changes showed a nesting pattern in space, with the spatially discrete sectors reflecting their mutational history (Fig. 3). A plausible branching pattern for the mycelium would reflect the phylogeny of nuclei over the growth by C1 from its point of origin.

**Figure 3.**
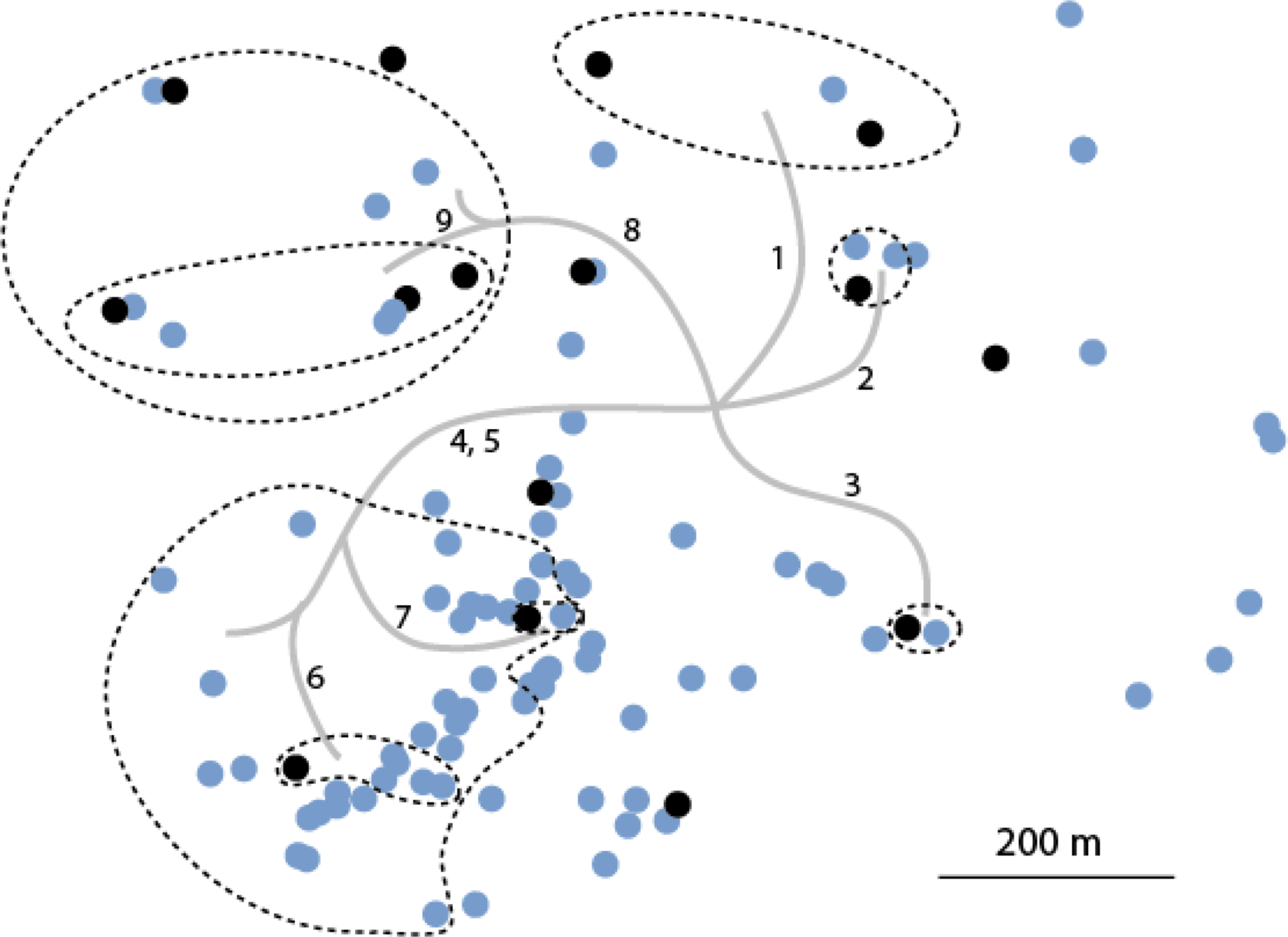
Variation at nine selected genomic sites mapped on all isolates of C1. Black dots represent the 15 Illumina sequenced isolates. The sites listed below were identified from the Illumina sequenced strains in Table S3. The sites were then PCR amplified and Sanger sequenced in other C1 isolates (table S4). Site variation is mapped to branches in the phylogenetic tree and spatial sectors encompassing derived genotypes are delineated with dashed lines. Site no. 1, scaffold 8, position 1611588; site no. 2, sc 24, pos 198035; site no. 3, sc 85, pos 12187; site no. 4 sc9, pos 1017767, site no. 5, sc 1, pos 1829026; site no. 6, sc 41, 509147; site no. 7, sc 23, pos 681486; site no. 8, sc 1, pos 1019568; site no. 9 sc 2, pos 3135125 (Supplementary Table 3).

Despite our ability to identify and map mutant alleles in the Armillaria mycelium, estimating the rate of mutation is problematic because we do not know the number of cell divisions intervening between any two isolates. We can estimate the number of genetic differences among the Illumina sequenced isolates experimentally, but the number of intervening cell divisions is inseparable from the mutation rate (no. differences = no. cell divisions times the mutation rate). Mutation rates, however, have been estimated in other organisms [19] and we can therefore assume an average mutation rate on the low end of the spectrum, 10^−10^ per base, per cell replication. (Note that mutation rate varies across the genome and among individuals [20].) With a genome of 100 mb, one mutation is expected every 100 cell divisions in a haploid genome and every 50 cell divisions in a diploid genome such as that of *A. gallica*. The pairs of strains were separated by an average of ca. 20 mutations, including those spatially separated by 1 km. In order to produce this number of mutations, the number of cell divisions would be restricted to ca. 1000 cell divisions over 1 km, or only one division for every 1 m of growth. If we assume a higher mutation rate, then the value of one cell division per meter of growth must increase even further. For example, a doubling of mutation rate to 2 ×10^10^, would restrict the number of cell divisions to 500, or only one division for every two m of growth over the 1 km span. Such a dearth of cell division is hard to reconcile with the length of cells within fungal hyphae, on the order of tens to hundreds of microns.

How might individuals of Armillaria protect themselves from mutation during cell division and growth? We see three possibilities, which are not mutually exclusive. First, the tips of rhizomorphs, which represent the inoculum potential for colonization, may remain relatively quiescent with respect to cell division, much like the apical meristem of plant roots [12, 21] and germline cells in mammals [22]. The rhizomorph tip, however, may be propelled forward by cell division and elongation behind the tip. In this way, the rhizomorph tip may minimize cell division even as it moves through its substrate. There is also precedent for avoidance of cell division in the shoots of plants in which axial meristems are derived from apical meristems with remarkably few intervening cell divisions [23]. A second possibility is that repair processes may have been driven to higher efficiency by natural selection up to the point where genetic drift negates any additional diminishing fitness benefit [24, 25]. Also, Armillaria exists in environments that may be of low mutagenic potential. UV radiation, for example, is low in such environments and damage such as pyrimidine dimer formation may be lower than in other environments. The third possibility is that the distribution of DNA strands after replication is asymmetric. Cells perpetuating the lineage might tend to receive old DNA while cells committed solely to local development and not to perpetuating the lineage would receive new DNA [26]. In Armillaria, this would mean that cells in the rhizomorph tip would retain the old DNA, whereas the subtending cells (committed to local, dead-end development) would receive the new DNA. In this way, the rhizomorph tips perpetuating the lineage would retain fewer mutations than cells committed to local differentiation. The extent to which each of these mechanisms may contribute to stability remains to be determined.

## 4. Conclusion

Here, we followed clonal evolution within cell lineages of a single fungal individual of *A. gallica* in a spatial context. This follows an earlier analysis of a much smaller individual of the same species in Ontario [12]. Our picture of clonal evolution in individuals of Armillaria closely parallels that in cancer progression within single individuals [27–30]. Cancer progression, however, is accompanied by extreme genomic instability [2], not necessarily due to loss of function in DNA repair processes, but rather to loss of control over DNA replication. The rate of replication increases to the extent that fidelity suffers and DNA damage accumulates rapidly. Evolution in cancer occurs on a time span shorter than the life span of the individual affected. Evolution occurs similarly in Armillaria individuals, but over a span of centuries and millennia, and is characterized by extreme genomic stability. The genomic stability of Armillaria and the underlying mechanisms allowing such stability may provide a useful counterpoint to cancer.

## Data Accessibility

The Illumina sequences for 15 collections of C1 are deposited as accession PRJNA393342 in the SRA at NCBI. All other data are in Supplementary Information as Tables S1, S2, S3, and S4.

## Authors’ contributions

J.N.B., J.B.A., and M.L.S. planned the study and did the field collecting. D.K. did the laboratory culturing and somatic compatibility tests, DNA extractions, PCR, Sanger sequencing, and preparation of DNA for Illumina sequencing. N.R. and H.W. did the bioinformatic analysis of the Illumina sequences and subsequent nuclear SNP discovery. J.B.A. did the analysis of mtDNA variation. J.N.B., J.B.A., and M.L.S. wrote the paper with input from D.K., N.R., and H.W.

## Supplementary Information

Bioinformatics S1

Table S1

Table S2

Table S3

Table S4

